# Response of methanogen community to elevation of cathode potentials in the presence of magnetite

**DOI:** 10.1101/2020.11.24.397190

**Authors:** Kailin Gao, Xin Wang, Junjie Huang, Xingxuan Xia, Yahai Lu

## Abstract

Electromethanogenesis refers to the process where methanogens utilize electrons derived from cathodes for the reduction of CO_2_ to CH_4_. Setting of low cathode potentials is essential for this process. In this study, we test if magnetite, an iron oxide mineral widespread in environment, can facilitate the adaption of methanogen community to the elevation of cathode potentials in electrochemical reactors. Two-chamber electrochemical reactors were constructed with inoculants obtained from a paddy field soil. We elevated cathode potentials stepwise from the initial −0.6 V vs standard hydrogen electrode (SHE) to −0.5 V and then to −0.4 V over the 120 days acclimation. Only weak current consumption and CH_4_ production were observed in the reactors without magnetite. But biocathodes were firmly developed and significant current consumption and CH_4_ production were recorded in the magnetite reactors. The robustness of electro-activity in the magnetite reactors was not affected with the elevation of cathode potentials from −0.6 V to −0.4 V. But, the current consumption and CH_4_ production were virtually halted in the reactors without magnetite when cathode potential was elevated to −0.4 V. Methanogens related to *Methanospirillum* were enriched on cathode surface of the magnetite reactors at −0.4 V, while *Methanosarcina* relatively dominated in the reactors without magnetite. *Methanobacterium* also increased in the magnetite reactors but stayed off electrodes in the culture medium at −0.4 V. Apparently, magnetite greatly facilitates the development of biocathodes, and it appears that with the aid of magnetite *Methanospirillum* spp. can adapt to high cathode potentials performing the efficient electromethanogenesis.

**IMPORTANCE:** Converting CO_2_ to CH_4_ through bioelectrochemistry is a promising approach for development of green energy biotechnology. This process however requires setting the low cathode potentials, which takes cost. In this study, we test if magnetite, a conductive iron mineral, can facilitate the adaption of methanogens to the elevation of cathode potentials. In the two-chamber reactors constructed using inoculants obtained from a paddy field soil, biocathodes were firmly developed in the presence of magnetite, whereas only weak electro-activity was observed in the reactors without magnetite. The elevation of cathode potentials did not affect the robustness of electro-activity in the magnetite reactors over the 120 days acclimation. *Methanospirillum* was identified as the key methanogens associated with cathode surface during the operation at relatively high potentials. The findings reported in this study shed a new light on the adaption of methanogen community to the elevated cathode potentials in the presence of magnetite.

## INTRODUCTION

Bioelectrochemical technology has been developed rapidly in recent decades with one of attracting applications being the conversion of CO_2_ to CH_4_ (1-3). Electromethanogenesis refers to the process where methanogens utilize electrons derived from cathodes for the reduction of CO_2_ to CH_4_ (4). Two plausible mechanisms have been proposed for electron transfer from cathodes to methanogens. One is the H_2_-mediated electron transfer, which assumes that H_2_ is electrochemically produced either abiotically or biotically that is then used by hydrogenotrophic methanogens for reduction of CO_2_ to CH_4_ (5, 6) and the other, though not fully proven, is the direct electron transfer from cathodes to methanogens (7-10).

Cathode potential is the critical factor controlling either chemical or biological H_2_ evolution. Theoretically, chemical H_2_ production from proton reduction can occur at −0.414 V (all potentials reported here are relative to standard hydrogen electrode, SHE), but in practice the cathode potential must be set substantially lower due to the overpotential in electrochemical operation (4, 11-13). Consequently, cathode potentials of −0.5 V to −0.8 V were usually applied in electromethanogenic reactors where H_2_-mediated electron transfer was assumed as the key process (4, 14-16). H_2_ evolution can occur through different bioelectrochemical mechanisms. For instances, it has been proposed that the high rate of hydrogen-mediated electromethanogenesis in *Methanococcus maripaludis* was due to the release of hydrogenases from living and dead cells, which attached to cathode surface and catalyzed H_2_ production (5). In mixed culture electrochemical systems, organisms other than methanogens may produce H_2_ by taking up cathode electrons directly or indirectly and then channel H_2_ to hydrogenotrophic methanogens for CO_2_ reduction to CH_4_ (6). In a defined coculture it was demonstrated that the Fe(0)-corroding sulfate-reducing strain IS4 performed direct cathode electron uptake during sulfate reduction with active H_2_ production at −0.4 V and −0.6 V, and H_2_ was consumed by *M. maripaludis* for CO_2_ reduction to CH_4_ (17).

Methane production from CO_2_ reduction can occur at a redox potential of −0.244 V under standard conditions. A few studies have suggested that some methanogens can operate the direct electron uptake from cathodes for catabolic metabolism, thus bypassing the H_2_-mediated processes. In this case it becomes less critical to apply a low potential as compared to the H_2_-dependent reactors. For instances, a marine-origin *Methanobacterium*-like strain IM1, which grew on iron specimen but hardly on H_2_, could sustain electromethanogenesis at −0.4 V, while the typical hydrogenotrophic *M. maripaludis* failed the operation at this potential (18). Similarly, *Methanosarcina barkeri*, a methanogen known to conduct direct interspecies electron transfer (DIET) with *Geobacter metallireducens*, can perform electromethanogenesis at −0.4 V (9, 10, 19). The hydrogenase-independent electron uptake by the *M. barkeri* mutant lacking hydrogenases has been observed at a potential of −0.484 V (8). Therefore, while H_2_-mediated electromethanogenesis generally requires low cathode potentials, H_2_-independent or DET-associated electromethanogenesis can work at relatively high potentials.

The performance of electromethanogenesis could be improved by introducing supplemental materials that are optimized for cathode electron transfer or that can efficiently catalyze H_2_ evolution at low potentials (20-22). Iron oxide minerals such as magnetite are common in soils (23-27). Recently, it has been demonstrated that the presence of magnetite nanoparticles (MNP) greatly promotes methanogenesis from oxidation of short-chain fatty acids in rice paddy soils and anaerobic sludges, and the stimulatory effect has been attributed to the facilitation of DIET (25, 27-30). Magnetite is a mixed-valent iron oxide mineral containing Fe(II) and Fe(III) in a ratio of 1:2. The edge-sharing of the octahedral Fe(II) and Fe(III) in magnetite facilitates the electron hopping or rapid electron exchange along the octahedral sublattice resulting in high electrical conductivity and redox activity (23, 31). It is tempting to explore the influence of MNP on electromethanogenesis.

Previous studies on electromethanogensis often collected inoculants from anaerobic digesters. Anaerobic digesters specialized in industry may limit the diversity of methanogens (1, 14, 15, 21, 22, 32), while natural environments such as paddy soil or wetlands can harbor a wider variety of methanogens (33) and hence offer an opportunity to screen methanogens capable of efficient electromethanogenesis. In the present study, electromethanogenic reactors were constructed using an inoculum obtained from a rice paddy soil. To test the effect of magnetite and explore how methanogens response to varying cathode potentials, the cathode potentials of reactors were elevated stepwise from −0.6 V to −0.5 V and finally to −0.4 V. Over four months of electromethanogenesis acclimation, we continuously monitored the CH_4_ production, the electrochemical performance and the microbial population dynamics. Systematic analysis of the combined data about the adaption of methanogens to the increasing cathode potential with and without magnetite sheds the new light on electro-active methanogens community originated from paddy soil.

## RESULTS

### Magnetite promoted electromethanogenesis

Three batches of reactors were constructed, namely the reactors with MNP (MNP reactors), the reactors without MNP (no-MNP reactors), and the reactors without inoculants but with MNP (the abiotic MNP control). All reactors were operated continuously except two breaks. In the initial phase, cathode potential was set at −0.6 V. Neither current consumption nor CH_4_ production were detected in the abiotic control. For the inoculated reactors, electromethanogenesis initiated after over a month acclimation. Magnetite exerted a strong influence. In the MNP reactors, rapid current consumption was detected at 33 d with concomitant sharp increase of CH_4_ production at 40 d (Fig. 2). The current consumption and CH_4_ production, however, were much lower in the no-MNP reactors where obvious electromethanogenesis occurred only after 50 d. When the current consumption in the MNP reactors did not increase further, the first break was applied. After the CV tests of cathodes and the exchange of catholytes with fresh culture medium, the circuits were reconnected and cathode potentials were elevated to −0.5 V for the second stage of operation. Methane production and current consumption occurred without delay in the MNP reactors, indicating that microbial populations retained on electrode surfaces (microbes in the catholytes were disposed during medium exchange) were well adapted to the electrochemical environment (Fig. 2). The current consumption and CH_4_ production also occurred in the no-MNP reactors, but the activity remained much lower as compared with the MNP reactors. The total CH_4_ accumulation was 22.68 mmol L^-1^ in the MNP reactors and 7.71 mmol CH_4_ L^-1^ in the no-MNP reactors, respectively, over the period at −0.5 V operation (Fig. 2a). When the current consumption stopped increasing again, the second break was implemented. After the CV tests of cathodes and culture medium exchange repeated, the cathode potentials were elevated to −0.4 V for the third stage of operation. The current consumption and CH_4_ production kept highly active in the MNP reactors. The total CH_4_ accumulation over 30 days operation actually exceeded those observed in the earlier stages at −0.6 V and −0.5 V. However, CH_4_ production and current consumption in the no-MNP reactors were virtually halted at −0.4 V (Fig. 2).

**Fig. 1.**
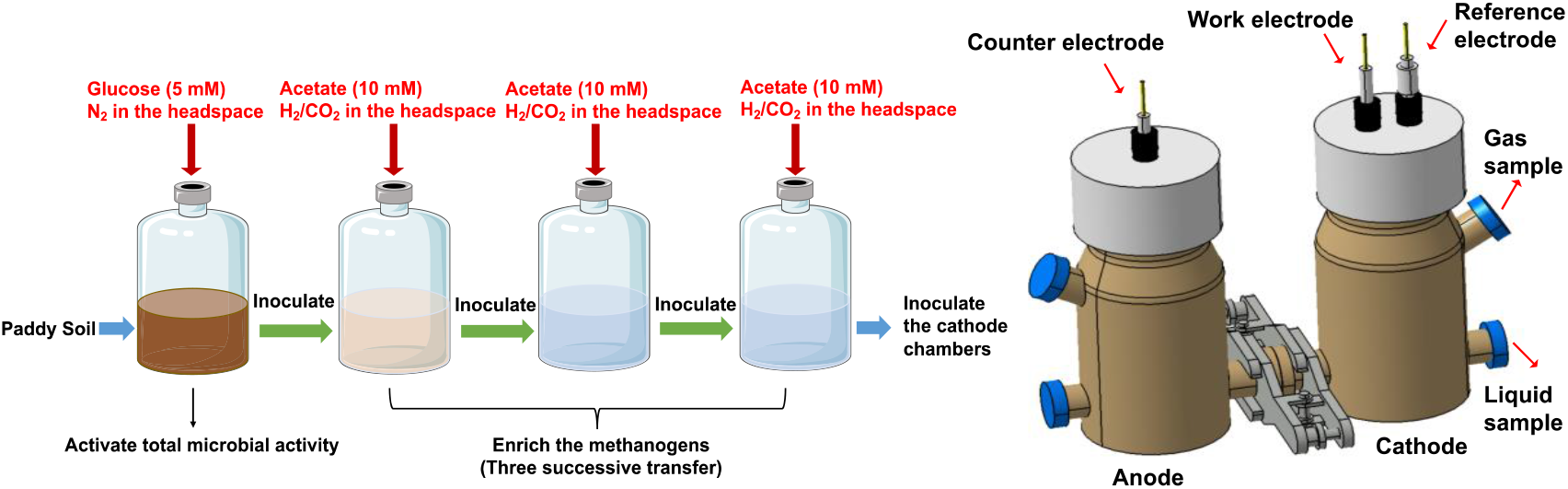
The schematic figure for enrichment cultivation of methanogens and the set of electrochemical reactors.

**Fig. 2.**
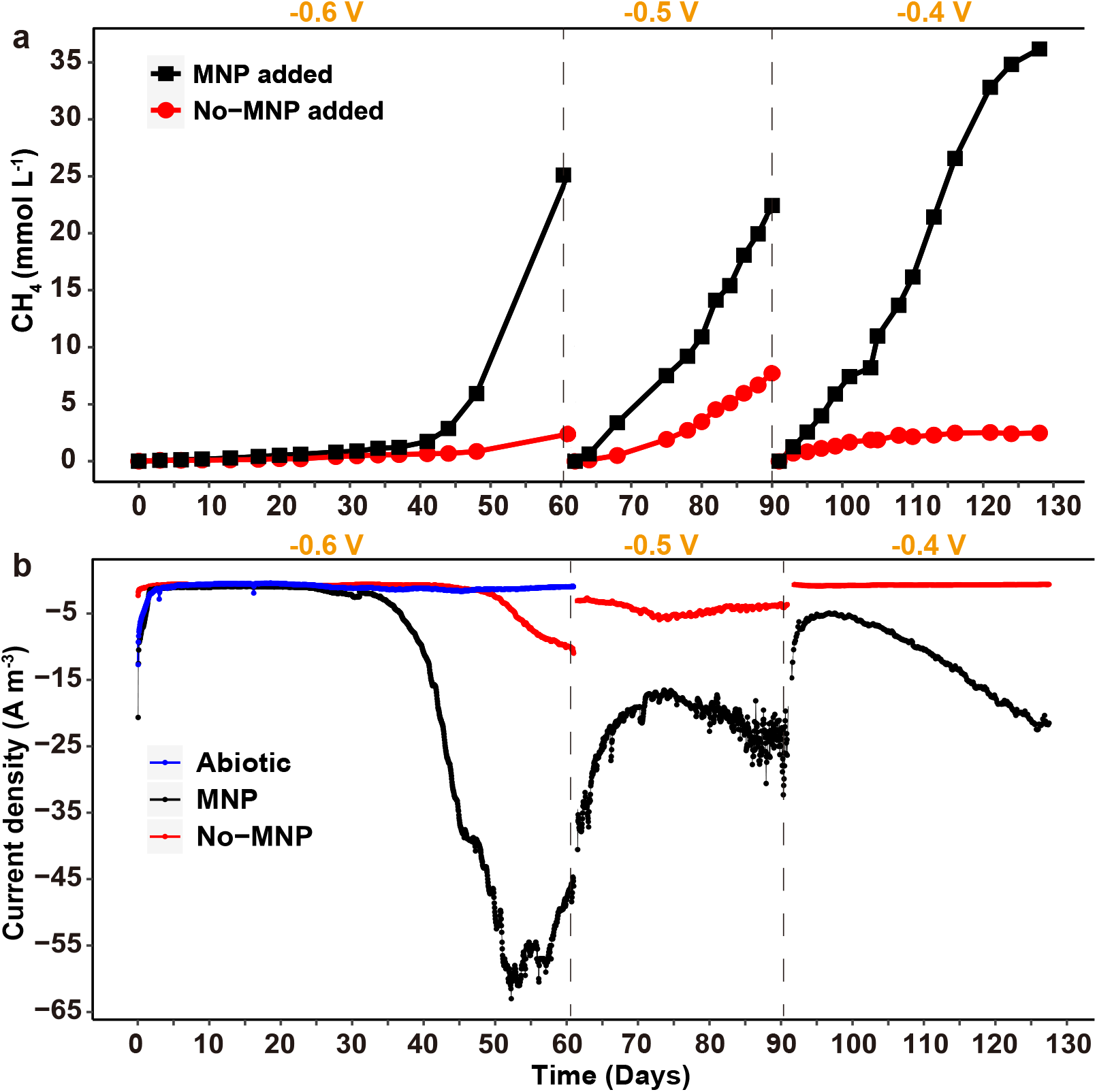
The production of CH_4_ and current density generation. The initial potential was set at −0.6 V and then was elevated to −0.5 V and −0.4 in two steps (yellow arrows). Data shown is a representative example of replicate or triplicate experiments (n=2 or 3), data for replicate or triplicate experiments could be found in Fig. S5 and S6

H_2_ accumulated to 0.02 mmol L^-1^ in the abiotic control during the initial operation at −0.6 V. H_2_ was occasionally detected in the MNP reactors at concentrations close to detection limit (0.01 mmol L^-1^). Otherwise H_2_ concentrations were below the detection limit in most cases, especially during the operations at −0.5 V and −0.4 V. To verify if addition of MNP influenced redox potentials (Eh) in culture medium, we measured Eh under open circuit conditions. The results showed no difference in the presence (−0.340 V) and absence (−0.338 V) of MNP.

At each break and at the end of experiment, CV tests were conducted to evaluate the catalytic activity of cathodes. At the break after −0.6 V operation, the no-MNP reactors revealed the lowest redox activity of cathodes, followed by the abiotic reactors while the highest redox activity was recorded for the MNP reactors (Fig. 3a). The activity of the abiotic reactors indicates that addition of MNP improved the conductivity of electrodes and electrolytes compared to the inoculated reactors without MNP. CV tests at the break after −0.5 V operation and at the end of experiment exhibited similar results and confirmed the significantly enhanced catabolic activity of cathodes in the MNP reactors compared with the no-MNP reactors (Fig. 3b, c). The voltammograms showed some distinct redox peaks and inflection points in the current profiles (Fig. 3a-c). For instances, at the break after −0.6 V operation, the current generation in the MNP reactors plateaued in the potential range from −0.23 V to −0.51 V, followed by an increase and then flattened again at −0.6 V. The inflection point occurred at −0.48 V for the MNP reactors at the break after −0.5 V operation and at −0.41 V after −0.4 V operation. These reflection points illustrated the robust development of biocathodes in the MNP reactors.

**Fig. 3.**
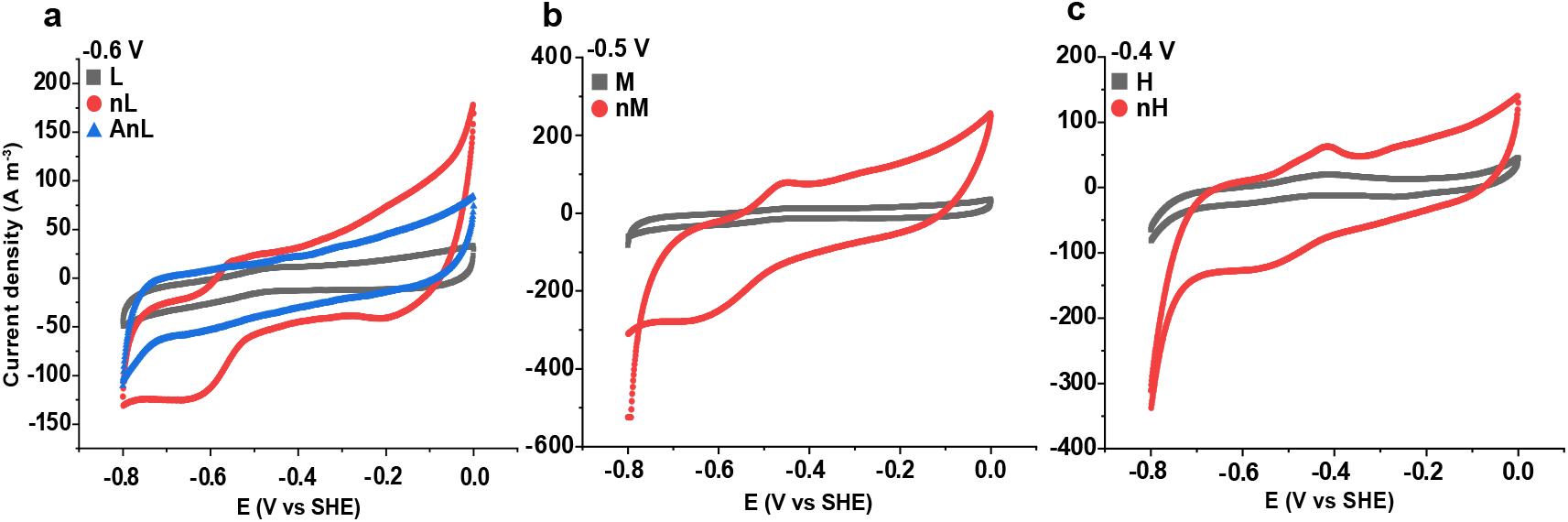
Cyclic voltammogram determined at the end of electromethanogenic operations under three successively elevated cathodic potentials (a, b, c separately). Red, MNP reactors; black, no-MNP reactors; blue, abiotic control reactors, which contained MNP but without inoculum. L, M, H denote −0.6 V, −0.5 V and −0.4 V; n, magnetite nanoparticles; A, abiotic control. Data shown is a representative example of replicate experiments (n=2 or 3), data for replicate experiments could be found in Fig. S7

### Response of archaeal and bacterial communities

Amplicon sequencing was used to analyze community compositions of archaea and bacteria attached on the cathode surfaces and living in the culture mediums (catholyte solution). The sequence summary for all archaea amplicons was given in Table S1. Archaea were composed exclusively of methanogen populations. Methanogens in the original inoculum obtained from rice paddy soil were dominated by *Methanospirillum* accounting for 90% of total archaeal sequences (Fig. 4a). The rest mainly affiliated to *Methanosarcina* and *Methanoregula*. Over the operation of electromethanogenesis, *Methanobacterium* replaced *Methanoregula*, arising as the third dominant methanogen (Fig. 4a; Fig. S1). The methanogen compositions in reactors were markedly influenced by the magnetite treatment, the potential elevation and the sample location (cathode surface vs catholyte medium).

**Fig. 4.**
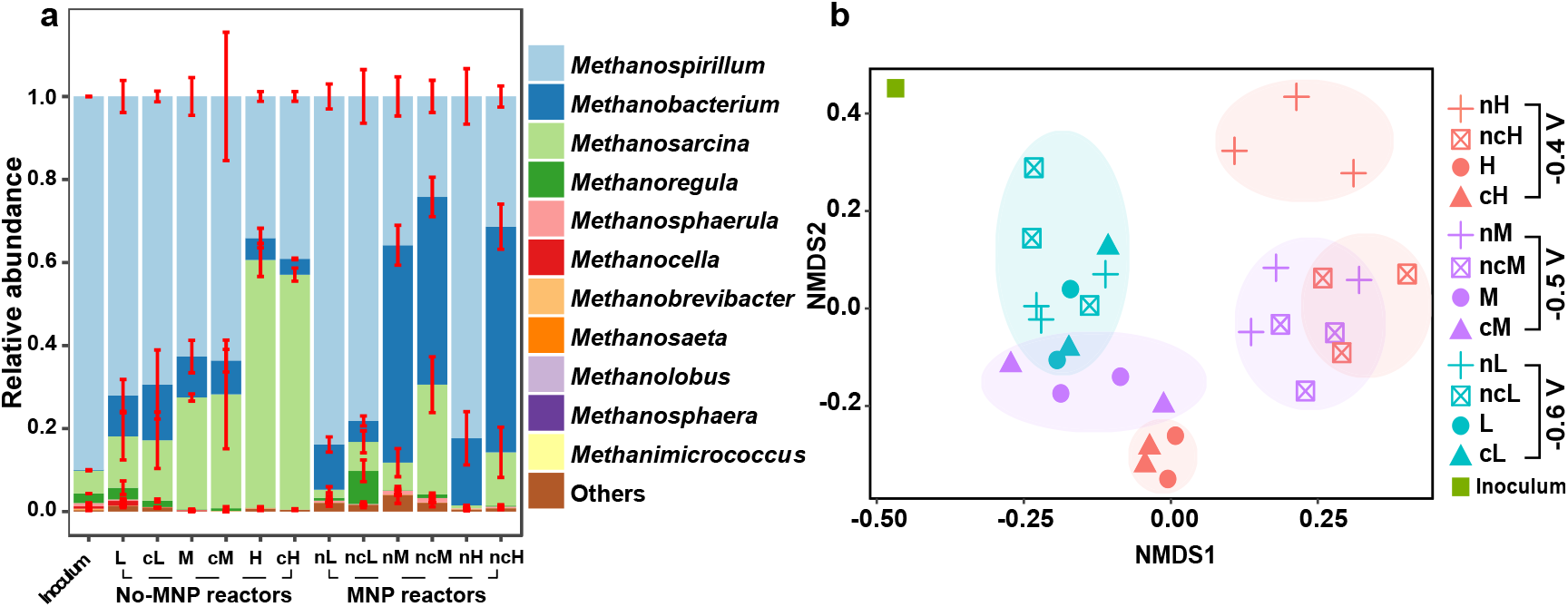
Composition dynamics of methanogens during the operation of electromethanogenesis. The relative abundance of methanogen community at genera level (a). Nonmetric multidimensional scale (NMDS) plot of methanogen community across samples (b). L, no-MNP biocathode at −0.6 V; nL, MNP biocathode at −0.6 V; cL, no-MNP catholyte at −0.6 V; ncL, MNP catholyte at −0.6 V; M, no-MNP biocathode at −0.5 V; nM, MNP biocathode at −0.5 V; cM, no-MNP catholyte at −0.5 V; ncM, MNP catholyte at −0.5 V; H, no-MNP biocathode at −0.4 V; nH, MNP biocathode at −0.4 V; cH, no-MNP catholyte at −0.4 V; ncH, MNP catholyte at −0.4 V. Error bars represent mean values ± one standard deviation, n=2 or 3

In the no-MNP reactors, the relative abundance of *Methanospirillum* and *Methanobacterium* decreased, while that of *Methanosarcina* increased from 15% to 60% with the elevation of cathode potentials from −0.6 V to −0.4 V (Fig. 4a). There was no significant difference in methanogen composition between the cathode surface and the culture medium (Fig. 4a). The MNP reactors showed a significantly different pattern. After the operation at −0.6 V, *Methanospirillum* remained dominant while *Methanobacterium* slightly increased compared with the original inoculum. When cathode potential was shifted to −0.5 V, *Methanobacterium* greatly increased and surpassed *Methanospirillum* both on cathode surface and in catholyte medium. *Methanosarcina* also increased relatively in the culture medium. When the cathode potential was further elevated to −0.4 V, *Methanospirillum* returned as the most dominant methanogen on cathode surface while *Methanobacterium* kept dominant only in catholyte medium (Fig. 4a). Apparently, *Methanospirillum* were selected against *Methanobacterium* and *Methanosarcina* on cathode surfaces at −0.4 V (Fig. 4a).

The shift of methanogen community was also illustrated by the nonmetric multidimensional scale (NMDS) analysis (Fig. 4b). Methanogen communities under electromethanogenesis were all distinct from original inoculum. At the break after −0.6 V operation, communities from all samples were clustered (green oval). At the break after −0.5 V operation, communities from the MNP and no-MNP reactors were separated (two purple ovals), while at the end after −0.4 V operation, a more significant divergence was revealed (three red ovals), with the separations not only depending on magnetite treatments but also on sample sites (cathode surface vs culture medium).

The phylogenetic relationship of top 10 archaeal OTUs was depicted in Fig. 5a. Clone_Mspi were closely related to *Methanospirillum psychrodurum* X-18 and *Methanospirillum lacunae* Ki8-1. Clone_Mba1 and clone_Mba2 were related to *Methanobacterium formicicum* MF and *Methanobacterium flexile* GH. Clone_Msar was related to *Methanosarcina horonobensis* HB-1 and *Methanosarcina mazei* DSM 2053. Given that *Methanospirillum* were predominant and selected by cathode surface in the MNP reactors at −0.4 V, we further analyzed the relative abundances of 15 OTUs affiliated to *Methanospirillum* (Fig. 5b). A significant shift was revealed for these *Methanospirillum* phylotypes. Clone_Mspi2 was dominated in the original inoculum. At the first break after −0.6 V operation, Clone_Mspi1, 4 and 14 increased, while clone_Mspi2 dropped. At the second break after −0.5 V operation, all the *Methanospirillum* OTUs declined. But at the end after −0.4 V operation, ten other OTUs including clone_Mspi3, 6-13 and 15 were substantially enriched (Fig. 5b). These results implied that clone_Mspi3, 6-13 and 15 instead of the dominant clone_Mspi2 in the original inoculum were enriched during electromethanogenesis at −0.4 V.

**Fig. 5.**
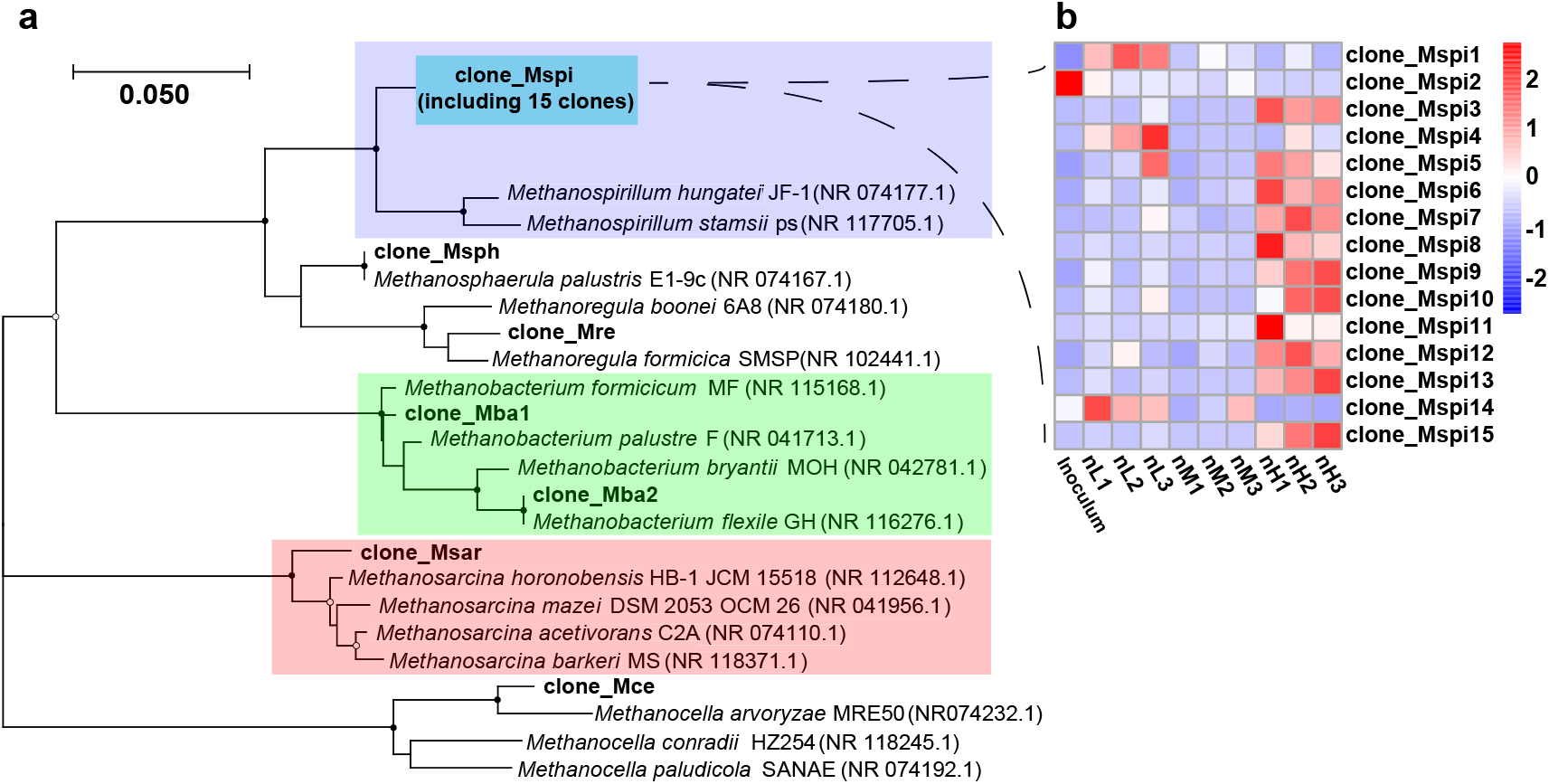
Neighbor-joining phylogenetic relationship of methanogens (a) and heatmap showing the change of phylotypes within *Methanospirillum* with the elevation of cathode potential (b). nL, MNP biocathode at −0.6 V; nM, MNP biocathode at −0.5 V; nH, MNP biocathode at −0.4 V

The sequence summary for the bacteria amplicons was listed in Table S2. Bacteria community consisted mainly of *Acetobacterium, Anaerolineaceae, Sulfuricurvum, Geobacter, Bacteroidales*, and *Desulfovibrio* (Fig. S2). *Acetobacterium* and *Anaerolineaceae*, the first and second most abundant bacteria across reactors, were more enriched in the MNP reactors compared to the no-MNP reactors (Fig. 6). By comparison, the other four bacteria mentioned above exhibited higher relative abundances in the no-MNP reactors than in the MNP reactors. The relative abundance of *Acetobacterium* however decreased sharply with the elevation of cathode potentials, from 54% to 10% in the MNP reactors and from 32.5% to 0.06% in the no-MNP reactors, respectively. Though relatively higher abundances of *Acetobacterium* on cathode surface than in catholyte medium of the MNP reactors at −0.6 V and −0.5 V, this difference was diminished at −0.4 V (Fig. 6). *Anaerolineaceae* showed the opposite tendency. Their relative abundances were low at −0.6 V and −0.5 V, but markedly increased with the elevation of cathode potentials to −0.4 V where they were the most dominant bacteria both on cathode surface and catholyte medium. *Actinobacteria* showed low relative abundance across all samples except on cathode surface in the MNP-reactors at −0.4 V where they became the second dominant bacteria after *Anaerolineaceae*. The NMDS analysis revealed that the samples from the MNP reactors at −0.6 and −0.5 V formed a cluster while those at −0.4 V formed another cluster (Fig. S4). The samples from the no-MNP reactors were separated into three clusters depending on cathode potentials and sample locations.

**Fig. 6.**
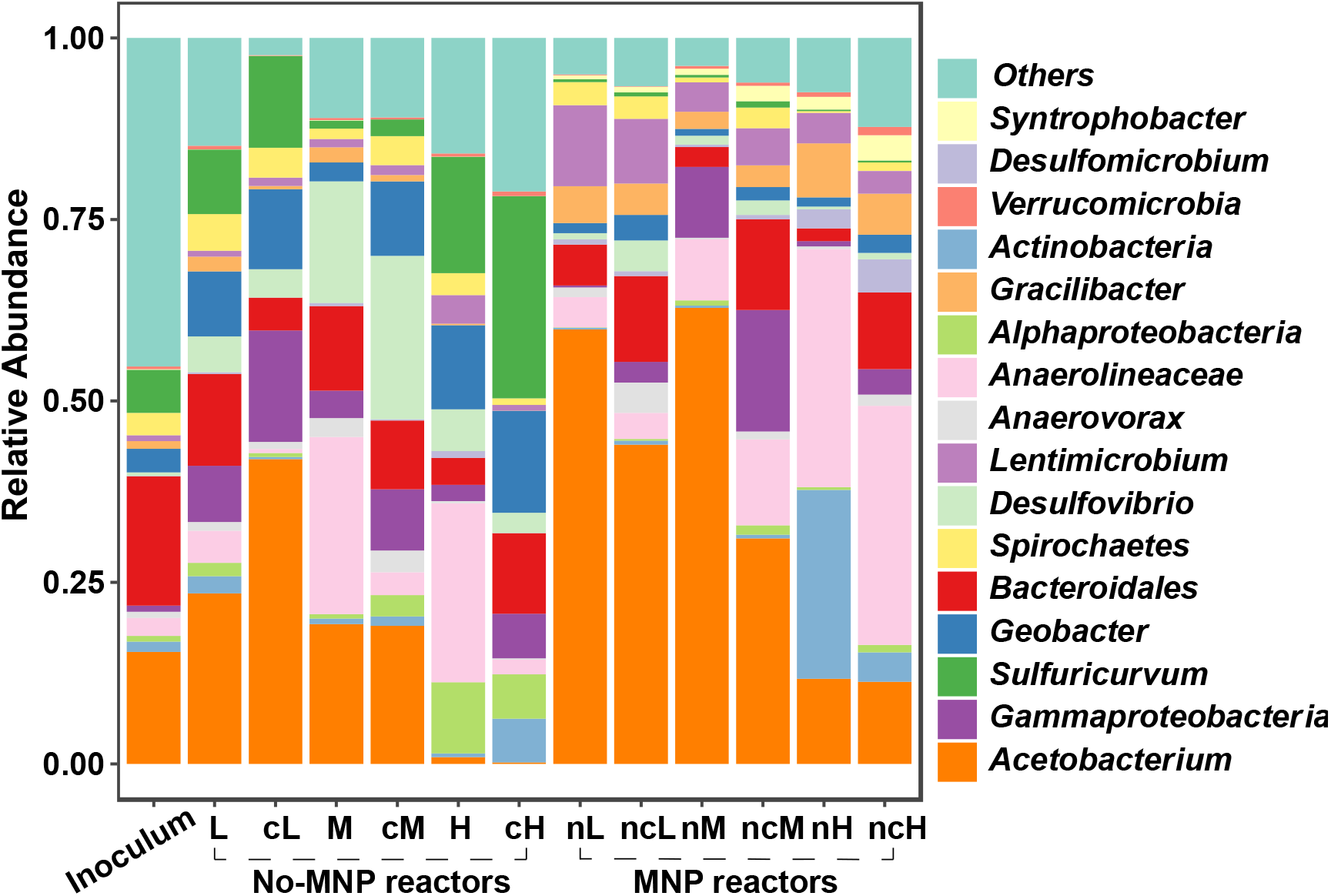
Bacterial community composition and relative abundance. Error bars represent mean values ± one standard deviation, n=2 or 3

## DISCUSSION

In the present study we demonstrated that electromethanogenesis derived from a paddy soil community was substantially promoted by magnetite during the 120 days operation with the elevation of cathode potentials from −0.6 V to −0.4 V (Fig. 2). Granular active carbon and zero-valent iron have been employed to accelerate the start-up of biocathodes and upgrade electromethanogenesis efficiency (20-22). One such a study however showed that electromethanogenesis was not affected by magnetite amendment in comparison with magnetite-free reactors (20). Thus, the stimulating effect appears dependent on reactor conditions and especially microbial identities in operations (34). Nevertheless, it has been demonstrated that in natural systems the syntrophic oxidation of short chain fatty acids and CH_4_ production were significantly stimulated by magnetite nanoparticles (25-27, 35). Magnetite was also shown to facilitate syntrophic interaction between a phototrophic bacterium *Rhodopseudomonas palustris* and an iron-reducing *Geobacter sulfurreducens* (36, 37). More recently, it was demonstrated that magnetite accelerated the aceticlastic methanogenesis by a pure culture *Methanosarcina mazei* zm-15 and its corresponding environment enrichment (38). The mechanisms are considered to be related to DIET in syntrophic interaction (25) or serving as an environmental battery facilitating electron transfer among microbes (37). In the present experiment, we lifted cathode potentials from the initial −0.6 V to the final −0.4 V over the 120 days acclimation. The electromethanogenesis remained robust in the presence of magnetite whereas the activity was substantially inhibited in the reactors without magnetite. Albeit mixture community in our reactors hindered the elucidation of exact mechanism, we assume that magnetite amendment facilitated the electron transfer from cathodes to methanogens either directly or indirectly.

Microbial communities in our reactors were markedly influenced by the magnetite treatment, the potential elevation and the sample location (Fig. 4, Fig. 6 and Fig. S3-S4). In the no-MNP reactors, the relative abundance of *Methanosarcina* gradually increased with the elevation of cathode potentials (Fig. 4a). *Methanosarcina barkeri* and *Methanosarcina mazei* have been demonstrated to be capable of DIET with *Geobacter metallireducens* or perform electromethanogenesis at −0.4 V (8-10, 19). It is possible that *Methanosarcina* spp. contributed to electromethanogenesis in the no-MNP reactors albeit their weak electrochemical performance. The relative abundance of *Methanosarcina*, however, substantially declined in the reactors with magnetite. Apparently, *Methanosarcina* did not respond to the stimulatory effect of magnetite or were less competitive during the development of biocathodes in the MNP reactors. *Methanobacterium* on the other hand significantly increased in the reactors with magnetite, especially at −0.5 V, reaching the relative abundance of about 50% on cathode surface as well as in catholyte medium. These methanogens however wane from cathode surface when cathode potentials were further elevated to −0.4 V. As the replacement, *Methanospirillum* were dominated on cathode surface at −0.4 V. The concomitant CH_4_ production and current consumption indicated that biocathodes were well developed in the MNP reactors (Fig. 2). The facts that the culture medium exchange at two breaks and the elevation of cathode potentials from −0.6 V to −0.4 V did not influence the electrochemical performance indicate that the biocathodes remained robust in the MNP reactors over the 120 days operation. This robustness of biocathode development was further illustrated with the CV tests at two breaks and at the end of experiment (Fig. 3). Accordingly, we assumed that microbial populations associated with cathode surface played the key role for electromethanogenesis in the MNP reactors. Many of previous studies have found that *Methanobacterium* were enriched in the cathode chambers of electrosynthesis reactors (13-16, 39). We show here that *Methanospirillum* were dominated on cathode surface at −0.4 V in the presence of magnetite (Fig. 4a). *Methanospirillum* have been underappreciated in the previous studies on electromethanogenesis. Recently, it has been reported that the archaellum of *Methanospirillum hungatei* is electrically conductive, making it an excellent candidate for the research of extracellular electron transfer in archaea and the application in electrochemical systems (40). It shall warrant a further investigation for whether the *Methanospirillum* in our reactors contain the electrically conductive archaellum, which facilitates the extracellular electron transfer in the biocathode ecosystems.

For the mixed culture reactors, the accompanying bacteria can play important roles in electromethanogenesis. Electro-acetogenesis refers to biological production of acetate from CO_2_ with electrons derived from cathodes (41, 42). *Acetobacterium* was the most dominant bacteria across all samples and were relatively enriched in the MNP reactors (Fig. 6), indicating their possible involvement in electro-acetogenesis. But the relative abundance of these acetogens significantly declined in the reactors at −0.4 V, suggesting they were not tolerant to the elevation of cathode potentials. The role of bacteria in the electromethanogenic reactors can be multifold. Firstly, acetogens produce acetate electrochemically which is then used by aceticlastic methanogens like *Methanosarcina* in the reactors. Second, some electrically-active bacteria may generate H_2_ by taking up electrons from electrodes and then channel H_2_ to hydrogenotrophic methanogens, forming syntrophic interaction (17). Third, it is plausible that DIET is established between the electrically-active bacteria and methanogens without the involvement of intermediate H_2_ production. Except *Acetobacterium*, some of potentially electrically-active organisms like *Geobacter*, *Sulfuricuvum* and *Desulfovibrio* were present as major bacteria members (Fig. S2). These organisms however exhibited relatively high abundances only in the no-MNP reactors, and thus did not play a significant role in the MNP reactors. *Anaerolineaceae* were the second dominant bacteria next to *Acetobacterium* (Fig. 6). These bacteria were significantly enriched in the MNP reactors at −0.4 V. *Anaerolineaceae* species have long filamentous structure (43, 44), and hence have an advantage of forming microbial aggregates or biofilms by attaching to electrode surface. Some *Anaerolineaceae* are known to ferment sugars and grow better in coculture with H_2_-consuming methanogens (44), implying their syntrophic life style. Therefore, it is plausible that *Anaerolineaceae* are involved in electromethanogenesis through forming syntrophic associations or improving methanogenic aggregate formation in the present experiment.

In summary, this study demonstrates that MNP greatly facilitates the adaption of methanogens in electrochemical reactors with the stepwise elevation of cathode potentials from −0.6 V to −0.4 V. *Methanospirillum* was identified as the key methanogens associated with cathode surface at −0.4 V. Given that the archaellum of *Methanospirillum hungatei* is known to be electrically conductive, it is worthwhile to further explore if *Methanospirillum* can perform direct electron transfer in the electrochemical reactors. Though a set potential of −0.4 V is considered to be sufficiently high to limit electrochemical H_2_ production under pure culture conditions (1, 8-10, 18), we are not able to dissect if the H_2_-mediated or H_2_-independent methanogenesis prevails in our reactors. Further research is necessary to determine if syntrophy is involved in electromethanogenesis, in which some bacteria may fetch electrons from electrodes forming H_2_ which is then utilized by the hydrogenotrophic *Methanospirillum* and *Methanobacterium*. It also remains unclear if the redox-active enzymes including hydrogenases are released from living and dead cells, which are then deposited on electrode surface facilitating H_2_-mediated processes (5, 45). These open questions await a further research in the future.

## MATERIALS AND METHODS

### Enrichment preparation for cathode inoculation

To focus on methanogen community and exclude carbon sources from soil, methanogenic enrichment from a rice paddy soil was prepared as below. The water-saturated soil samples were collected from a paddy field located in the northeastern China close to Heihe city of Heilongjiang province, China (127.36°E, 49.90°N). Four successive transfers of anaerobic incubation were conducted. For the first transfer, fresh soil samples were suspended in autoclaved degassed water at a water to soil ratio of 5:1 (soil mass in dry weight). Aliquots (50 ml) of the homogenized soil slurry were then dispensed into 125 ml sterile serum bottles. Glucose was added into bottles at a final concentration of 5 mM in slurries. Headspace of serum bottles were flushed with N_2_ thoroughly. All the bottles were sealed with butyl stoppers and aluminum crimp caps, and put in the dark at room temperature (27 °C). When the rate of daily CH_4_ production reached to the quasi-steady state, the soil slurries were used as inoculum (5% v/v) for the next three transfers where acetate (10 mM) was spiked into 50 ml HEPES-buffered (30 mM, pH 7) culture medium and the headspace of serum bottles were flushed with H_2_/CO_2_ (80:20; v/v). Both acetate and H_2_/CO_2_ served as the carbon and energy source for methanogen enrichment (Fig. 1). The basal medium consisted of MgCl_2_·6H_2_O (0.4 g/L), CaCl_2_·2H_2_O (0.1 g/L), NH_4_Cl (0.1 g/L), KH_2_PO_4_ (0.2 g/L), KCl (0.5 g/L). Supplements of vitamin, and trace element solutions were applied. Resazurin (46) and cysteine were omitted to avoid the possible effect of electron shuttle molecules (47). All the bottles for the enrichment cultivation were sealed and put in the dark at room temperature (27 °C). When acetate was used up in the medium, the enrichment cultures were used to inoculate (10% v/v) the cathode chambers of bioelectrochemical reactors. Abiotic control reactors were established without inoculation.

### Setup of bioelectrochemical systems

The bioelectrochemical reactors consisted of two chambered borosilicate gastight H-type microbial electrolysis cells, in which the 250 ml anode and cathode chambers were separated by a Nafion 117 proton exchange membrane (surface area 5 cm^2^, DuPont, Wilmington, DE, USA) (Fig. 1). Prior to use, membranes were successively treated one hour each with H_2_O_2_ (3.5 wt.%), distilled water, H_2_SO_4_ (5 wt.%), and distilled water. In each chamber, there are an upper and a lower openings, which were sealed with butyl stoppers and aluminum crimp caps. The upper openings were used to collect gas samples (Fig. 1). Carbon fiber brush (volume 20 cm^3^) acted as the work electrode in the cathode chambers and platinum foil was used as the counter electrode in the anode chambers. An Ag/AgCl reference electrode (+0.2046 V vs SHE, 25°C) was placed in the cathode chamber as a reference electrode. Magnetite nanoparticles (MNP) was synthesized via a conventional aqueous co-precipitation method (48) as described previously (27).

### Bioelectrochemical measurements and cathode potential elevation

Nitrogen was flushed up to one hour to make the anaerobic condition. After autoclaving, the anaerobic basal medium was added into two chambers of reactors. The composition of basal medium was same as the enrichment cultivation but without the addition of acetate. 125 ml medium was dispensed into anode chambers. And 112.5 ml of basal medium and 12.5 ml of inoculum were dispensed into cathode chambers to bring the total volume to 125 ml. The abiotic control was prepared with addition of 125 ml same medium in both chambers without inoculation. The headspace of each reactor had a volume of 125 ml and was flushed with N_2_/CO_2_ (80:20; v/v) thoroughly. CO_2_ was the sole carbon source in reactors. Reactors were put in the dark at room temperature (27°C). The experiment was divided into three groups. For the first group, MNP was added into the cathode chambers with a final concentration of 10 mM in Fe atom. The second group was prepared similarly but without MNP in chambers. The third group was set up as abiotic control with addition of MNP in cathodes but without inoculation of enrichment. All reactors were connected into the eight channel potentiostat (CHI 1000C, CH Instruments Inc., Shanghai, China) which recorded current every 100 seconds automatically. The reactors were operated in a continuous mode except two breaks for the shifting of cathode potential. The initial cathode potential was set at −0.6 V vs standard hydrogen electrode (SHE, all potentials below, unless specified, were all versus standard hydrogen electrode). When the current density of MNP bioreactors showed no further change or started falling, all the reactors were disconnected from the potentiostat for the first break. The cyclic voltammetry (CV) was conducted immediately with a scan rate of 5 mV/s and scan range of −0.8 V to 0 V vs SHE to characterize biocathodes. Microbial samples were also collected immediately from the cathodes and catholyte medium (see details below). The reactors were then reconnected to the potentiostat with the cathode potential elevated to −0.5 V and operated until the current density of MNP bioreactors started falling again. The second break was then applied for CV measurement and microbial sampling. Finally, the cathode potential was raised to −0.4 V for the last round of electrochemical operation. At the end, the last CV tests and microbial sampling were performed again.

During each break, the chambers were rinsed thoroughly to remove remaining microbes and magnetite, then replenished with the fresh medium. Cathode chambers were inoculated by 10% (v/v) of the used medium and the remaining used medium was discarded. The electrodes with biofilm remained unchanged. Magnetite nanoparticles (MNP) were re-supplemented in the MNP reactors with the same concentration as before. Microbial sampling and medium refreshing were performed in an anaerobic glove box. There were nine reactors in total at the beginning, i.e. three each for MNP bio-reactors, no-MNP bio-reactors and MNP abiotic reactors. But during the first-round operation at −0.6 V, one of the no-MNP biotic reactors failed to produce CH_4_ after 50 days and hence this reactor was suspended. Thus, there were eight functioning reactors in operation throughout the experiment.

### Chemical analyses

Methane and hydrogen were monitored throughout experiments. Gas samples (200 μL) were regularly taken from the headspace of cathode chambers with a Pressure-Lok precision analytical syringe (Bation Rouge, LA, USA). The concentration of CH_4_ and H_2_ were analyzed using GC-7890B gas chromatograph (Agilent Technologies, Santa Clara, CA, USA) equipped with flame ionization detector. The detection limits are 5 Pa for both CH_4_ and H_2_. Every time after gas sampling, chambers were shaken vigorously for one minute.

### Microbial community analysis

At the end of CV tests, an aliquot of culture medium was collected, while small pieces of carbon brushes were cut off from electrodes (without influencing the electrode integrity). Carbon brush samples were washed twice with sterile demineralized water to remove the loosely attached microbial cells. Microbial DNA were extracted from both culture medium and carbon brush samples. FastDNA™ SPIN Kit (MP Biomedicals, Irvine, CA, USA) was used for DNA extraction according to the manufacturer’s instructions. Archaeal DNA was amplified using the 16S rRNA gene primers 1106F (TTWAGTCAGGCAACGAGC) and 1378R (TGTGCAAGGAGCAGGGAC) (49). The bacterial DNA was amplified using the 16S rRNA gene primers 515F (GTGCCAGCMGCCGCGGTAA) and 806R (GGACTACHVGGGTWTCTAAT). Amplicon sequencing was completed by Novogene (Beijing, China) using Ion S5XL platform. About 80000 reads were obtained for each sample and were clustered into operational taxonomic units (OTUs) with a similarity threshold of 97%. Nonmetric multidimensional scale (NMDS) was conducted using R and vegan community ecology package (50). The closest matching sequences were identified by searching with the BLAST program in the NCBI database (51). Neighbor-joining phylogenetic trees were constructed using MEGA-X and presented with ggtree package in R (52, 53). The nucleotide sequences generated from this study has been deposited in SAR database under the accession numbers of SAMN13136352-SAMN13136413.

## ACKNOWLEDGEMENTS

This study was financially supported by the National Natural Science Foundation of China (No. 41630857; 91951206).

## SUPPLEMENTAL MATERIAL

Sequence summary for amplicon sequencing of microbes, total abundance of methanogens and bacteria in electromethanogenic reactors, relative abundance of bacterial community, nonmetric multidimensional scale (NMDS) plot of bacteria, CH_4_ production, current density generation and CV tests in replicate or triplicate reactors.

